# Comparative Evaluation of Micronaut-AM and CLSI broth micro-dilution method for antifungal susceptibility testing of Aspergillus species against four commonly used antifungals

**DOI:** 10.1101/811901

**Authors:** Ali Nuh, Newara Ramadan, Silke Schelenz, Darius Armstrong-James

**Affiliations:** Department of Microbiology, Laboratory Medicine, Royal Brompton and Harefield NHS foundation trust, London; Department of Microbiology, Kings College Hospital, London

**Keywords:** *Aspergillus species*, Susceptibility Testing, Micronaut, Colorimetric Method, alamarBlue

## Abstract

The aim of this study was to evaluate a colorimetric method, Micronaut-AM, for determining susceptibility testing of anidulafungin, amphotericin, voriconazole and itraconazole by comparing the Minimum Inhibitory (Effective) Concentrations (MICs/MECs) obtained by this method to those generated by the reference Clinical Laboratory Standard Investigation (CLSI) method. 78 clinical isolates of *Aspergillus species*, nine of them azole-resistant, were tested against above antifungals. *A fumigatus* ATCC 204305 was used as a reference strain and test was performed in accordance with slightly modified yeast susceptibility testing instruction of the manufacture; conidia suspension inoculum and alamarBlue concentration were optimised. These same isolates were referred to Bristol Mycology reference laboratory and tested by CLSI method. The MICs and MECs generated by the two methods were compared using concordance analysis.

Micronaut-AM (MN) showed significant concordance (P< 0.0001) with CLSI method and overall agreement was high (≥ 90%). In addition, Micronaut-AM produced echinocandin MECs results within 18-24h incubation time and reliably detected azole resistant isolates. Essential agreement (within 2 log2 dilution of median) between MN and CLSI reference laboratory method was 99 % for anidulafungin, 100 % for amphotericin; 90% for voriconazole and 87 % for itraconazole. Categorical agreement for anidulafungin, amphotericin B, voriconazole and itraconazole were 100%, 96%, 97% and 99% respectively.

Micronaut-AM showed very good agreement with the reference broth micro-dilution method results for all antifungal agents tested and was able to detect azole resistance. This colorimetric method is very promising and appears to be a suitable alternative susceptibility testing method to labour intensive broth microdilution method for *Aspergillus species*.

## Introduction

Aspergillosis alludes to a spectrum of fungal diseases which are caused by *Aspergillus species*; these include invasive aspergillosis, allergic bronchopulmonary aspergillosis and chronic pulmonary aspergillosis.^1^ Invasive aspergillosis is a fatal and difficult to treat infection that mainly affects immunocompromised and critically ill patients. The mortality rate of this infection is very high and successful clinical outcome depends on early start of systemic antifungal treatment, which in turn might be optimised by the prompt availability of an in vitro susceptibility result.^2,3^ Susceptibility testing of all significant isolates of *Aspergillus species* is recommended by British Society of Medical Mycology as the best practice.^4^ The availability of in vitro antifungal susceptibility is becoming even more critical considering emerging and increasing resistance of *Aspergillus species* to mainstay antifungal agents, which in turn increases mortality; for instance, patients with azole resistant invasive aspergillosis have higher mortality rate compared with those with azole sensitive invasive aspergillosis.^5,6^ Both the European Committee on Antimicrobial Susceptibility Testing (EUCAST) and the Clinical and Laboratory Standards Institute (CLSI) recommend broth micro-dilution method (BMD) for filamentous fungi susceptibility testing, including Aspergillus species.^7,8^

Microbroth dilution testing is labour intensive and needs expertise which may not be available at the local clinical microbiology laboratories and its introduction into routine use remains a challenge. Mould susceptibility testing, especially *Aspergillus species*, was available only 17% of 154 laboratories surveyed in 2017 in the UK.^9^ In light of this, a few commercial susceptibility testing methods have been marketed by different companies. Among these a broth micro-dilution-based colorimetric method, Sensititre YeastOne (YO), has been widely evaluated and shown the best correlation with microdilution reference method.^10–13^ Commercially available colorimetric methods are alternatives to reference methods and offer many practical advantages including ease of performing test and less subjective Minimum Inhibitory Concentration (MIC) determination. These methods require only the addition of conidia containing media and MIC end point reading is facilitated using the metabolic dye, alamarBlue.

Despite the reported high correlation of YO with BMD, one drawback of this method is that micro-titre plates come with alamarBlue already added, therefore it is not possible to optimise this growth indicator. AlamarBlue optimisation is very important as different filamentous fungi has different metabolic rate and isolates from different patient populations may vary. Lack of this optimisation may explain high amphotericin MICs in *A fumigatus* isolated from cystic fibrosis patients reported by Dunne et al.^14^ and low echinocandin concordance by Siopi et al.^15^ 12 out 56 *A fumigatus* isolates tested by Dunne et al. had amphotericin MIC of 2mg/L. Interestingly, we obtained similar results when locally validating YO and a significant number of isolates in our collection were from cystic fibrosis patients (data not published). Siopi et al. suggested that YO AlmarBlue concentration may not be suitable for fast growing Aspergillus species such as *A flavus* and that further optimisation of growth indicator is warranted for each species. Micronaut-AM (Merlin Diagnostika GmbH) is a commercially available colorimetric method similar to YO except it supplies alamarBlue separately as supplement. This EUCAST-based broth microdilution method has been mainly developed for yeast susceptibility testing and consists of dried antifungal agents in micro titre plate, RPMI 1640 growth medium and growth indicator, AlamarBlue, which is supplied separately as a supplement for the improvement of visual reading of MIC end points.

The aim of the current study was to evaluate Micronaut-AM for *Aspergillus species* susceptibility testing by optimising alamarBlue concentration; inoculum and incubation time and in turn increase the repertoire of colorimetric methods available for routine use. To our knowledge there is no published work on the use of this method for filamentous fungi including *Aspergillus species* and no instruction was available for filamentous fungi susceptibility testing.

## Material and Methods

A total of 78 clinical isolates of Aspergillus species (46 *A. fumigatus*; 14 *A. flavus*; 10 *A terreus* and *10 A nidulans*), nine of them azole resistant, from the private collection of the department of medical microbiology of RBHT (mainly isolated from respiratory samples) were tested. This collection of clinical Aspergillus species isolates was also routinely referred to Bristol Mycology reference laboratory, where they were tested by CLSI microdilution method. MICs generated by MN were compared with CLSI MIC results.

### Micronaut-AM method

Conidial inoculum suspension was prepared from cultures grown on potato dextrose agar (Oxoid Microbiology products, UK) at 35°C for 5 days. Conidia were harvested with cotton swabs and suspended in 0.05% Tween saline solution. The conidial suspension was adjusted to 0.5 McFarland Standard by nephelometer (Sensititre^R^ Nephelometer^R^) and further diluted to 1:56 in RPMI broth medium supplemented with alamarBlue (0.009% v/v), which was used as a working solution. This was achieved by adding 200μL of suspension to 11ml of RPMI-Broth (Merlin Diagnostica GMBH). Then 100μL of working solution was dispensed into each micro-titre plate well with a multi-channel pipette and panels were covered with adhesive seal strip and incubated at 35°C for 48 hours. Plates were inspected and read at 18-24h and 48h. *A fumigatus* ATCC 204305 was used as a reference strain and was tested in each experiment.

Minimum Inhibitory Concentrations (MICs) for Amphotericin, Itraconazole and Voriconazole were interpreted as the lowest drug concentration that showed complete fungal inhibition as indicated by blue colour. For anidulafungin, the Minimum Effective Concentration (MEC) was determined as the first purple well after 18-24h incubation and confirmed by viewing aberrant growth (abnormal, short and branched hyphal clusters) by using inverted microscope. The first purple well at 18-24h was marked and checked again at after 48h incubation. Isolates giving discrepant MIC/MEC results from CLSI results were repeated.

As for reproducibility studies, four isolates of *A fumigatus, A flavus, A terreus* and *A nidulans* were tested on three different days and read by three different observers. MICS and MECs were interpreted as above.

### Data analysis

High off-scale MICs and MECs were rounded to the next higher concentration, whilst the low off-scale values were left unchanged. Median, geometric mean, range and essential percent agreement were calculated for each species and drug (table 1). MICs and MECs values were transformed into Log2 to normalise the data and percent agreement was calculated. Essential agreement was defined as the proportions of MICs/MECs that fall within two-log2 dilution of the median, whereas categorical agreement was defined as the proportions of results that were classifies as either sensitive or resistant, according to epidemiological cut-off, and agree in the two methods. Percent essential agreement and categorical agreement were determined for each drug using Kappa interrater agreement. Statistical analysis was carried out by using STATA^R^ Software (StataCorp. 2017). For reproducibility studies, essential agreement (defined also as above) between experiments and observers was calculated. For echinocandin only anidulafungin was included in the data analysis since no CLSI data were available for other drugs.

**Table 1:** MICs (mg/L) rages, median and geometric (GM) mean for Anidulafungin (Anid), Amphotericin (Aph), Itraconazole (Itr) and Voriconazole (Vor) determined by Micronaut-AM and CLSI methods.

| Isolate (No tested) |  | Micronaut-AM Method |  |  |  | CLSI Method |  |  |  |
| --- | --- | --- | --- | --- | --- | --- | --- | --- | --- |
|  |  | Anid | Aph | Vor | Itr | Anid | Aph | Vor | Itr |
| *A. flavus*<br>(14) | Range | 0.002-0.015 | 0.500-2.00 | 0.0625-0.500 | 0.031-0.500 | 0.002-0.015 | 0.500-2.00 | 0.250-1.00 | 0.0625-0.500 |
|  | Median | 0.002 | 0.500 | 0.250 | 0.063 | 0.002 | 1.000 | 0.500 | 0.125 |
|  | GM | 0.003 | 0.743 | 0.226 | 0.097 | 0.002 | 1.000 | 0.431 | 0.152 |
| *A. fumigatus*<br>(46) | Range | 0.002-0.015 | 0.250-1.00 | 0.031-1.00 | 0.031-8.00 | 0.002-0.015 | 0.250-1.00 | 0.125-4.00 | 0.031-32.00 |
|  | Median | 0.002 | 0.500 | 0.125 | 0.091 | 0.002 | 0.500 | 0.250 | 0.125 |
|  | GM | 0.002 | 0.478 | 0.135 | 0.155 | 0.002 | 0.457 | 0.348 | 0.299 |
| *A. nidulans*<br>(8) | Range | 0.0020-0.015 | 0.250-2.00 | 0.0625-0.500 | 0.031-0.250 | 0.002-0.015 | 0.250-2.00 | 0.0625-1.000 | 0.0625-1.00 |
|  | Median | 0.015 | 0.500 | 0.125 | 0.094 | 0.002 | 0.500 | 0.250 | 0.500 |
|  | GM | 0.013 | 0.545 | 0.136 | 0.096 | 0.003 | 0.545 | 0.296 | 0.386 |
| *A. terreus*<br>(10) | Range | 0.002-0.015 | 0.500-1.00 | 0.0125-2.00 | 0.031-4.00 | 0.002-0.015 | 0.500-2.00 | 0.250-4.00 | 0.031-1.00 |
|  | Median | 0.002 | 0.500 | 0.094 | 0.031 | 0.002 | 1.000 | 0.500 | 0.094 |
|  | GM | 0.002 | 0.616 | 0.140 | 0.094 | 0.002 | 0.933 | 0.489 | 0.125 |
| Overall (78) | Median | 0.002 | 0.500 | 0.125 | 0.0625 | 0.002 | 0.500 | 0.250 | 0.250 |
|  | GM | 0.0028 | 0.542 | 0.149 | 0.127 | 0.0022 | 0.587 | 0.372 | 0.243 |

A very major discrepancy was defined when Micronaut categorised an isolate as sensitive and CLSI categorised as resistant. A major discrepancy was defined when Micronaut categorised an isolate as resistant and CLSI categorised as sensitive.

## Results

MICs for the quality control strain were within the expected range and MN method reliably detected eight out of nine azole resistant isolates; the remaining isolate was classified as wild type by MN in contrast to CLSI and no resistance was detected when inoculated onto VIPcheck™ system for azole resistant screening. This isolate has an MIC of 0.25mg/L in MN method compared with CLSI MIC of 1.5mg/L. 48h was sufficient incubation time and all isolates tested were able to change alamarBlue to its pink derivative and negative and positive control in MN consistently showed the expected colour and results and no discrepancies were observed. Echinocandin wells were all pink at 48h incubation, however, MEC end points were consistently indicated by the first purple well at 18-24h incubation, which was confirmed by viewing aberrant hyphal growth; the wells showing aberrant growth did not change after 48h incubation.

MICs ranges, median and geometric mean for both MN and CLSI methods are presented in table 1. Generally, both methods gave close and comparable median and geometric mean for all isolates, except that amphotericin median and geometric mean was slightly higher in CLSI for *A flavus* and *A terreus*. In addition, overall itraconazole and voriconazole median MICs were slightly lower in MN compared with CLSI. Itraconazole MN median MIC (0.0625mg/L) was two dilutions lower than that of CLSI (0.25mg/L), whereas voriconazole MN median MIC (0.125mg/L) was just one dilution lower compared with CLSI results (0.25mg/L). Median MEC of anidulafungin and median MIC amphotericin were the same for both methods for all isolates tested; 0.002mg/L and 0.50mg/L respectively.

Both essential and categorical agreements between the two methods were very high (≥ 90%) and few major discrepancies (MD) and very major discrepancies (VMD) were observed. Overall essential agreement between Micronaut-AM generated results and the reference CLSI result, as defined by the proportion of MICs/MECs that fall within two-doubling dilution of the median, was 99% for anidulafungin, 100% for amphotericin, 90% for voriconazole and 87% for itraconazole (table 2). MN also showed excellent percent categorical agreement with CLSI results. Percent categorical agreements were 100%, 96%, 97% and 99% for anidulafungin, amphotericin B, voriconazole and itraconazole respectively (table 2).

**Table 2:**
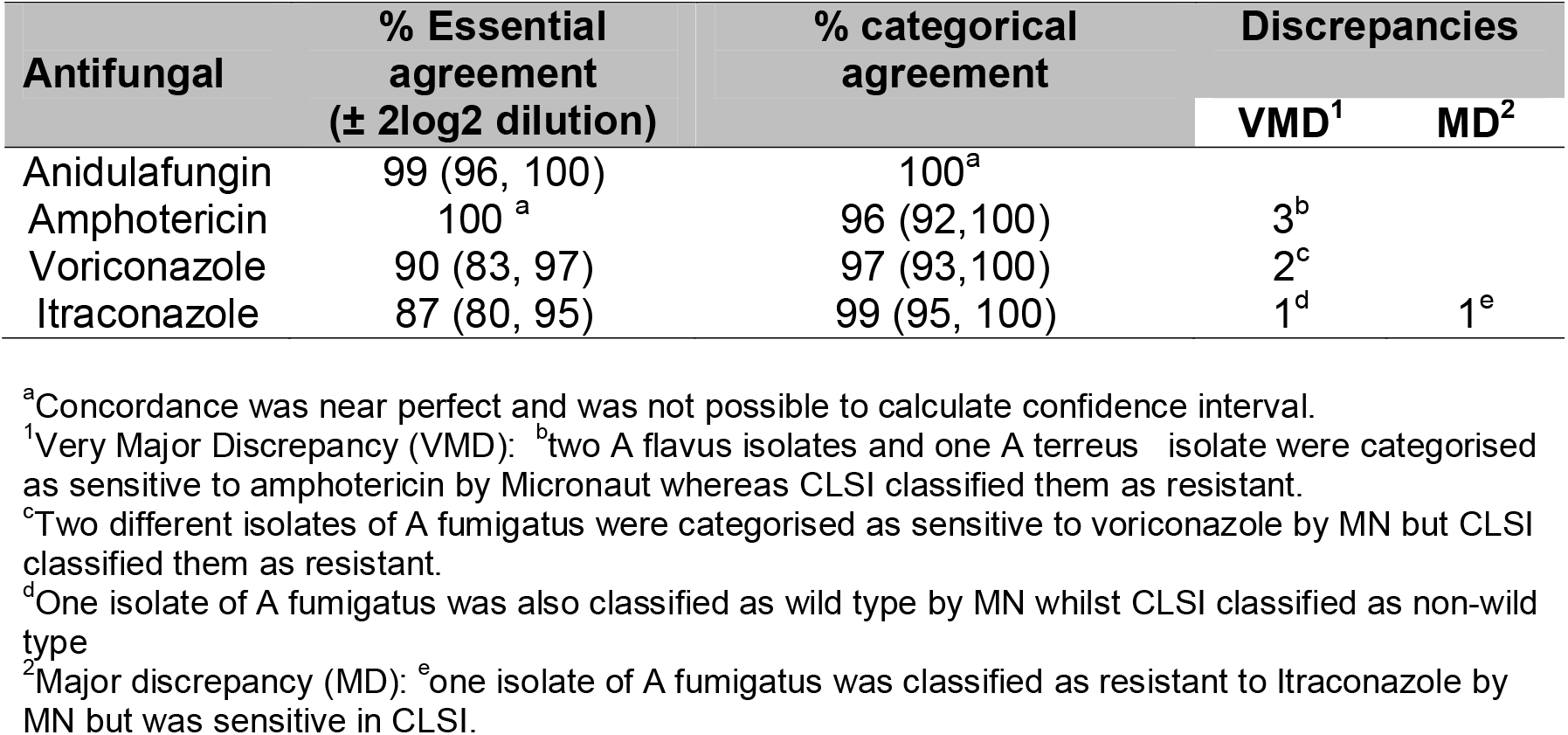
Overall percent essential agreement (with 95% Confidence Interval) and categorical agreement between Micronaut-AM and CLSI methods for all 78 Aspergillus isolates.

| Antifungal | % Essential agreement<br>(± 2log2 dilution) | % categorical agreement | Discrepancies | |
| --- | --- | --- | --- | --- |
|  |  |  | VMD<sup>1</sup> | MD<sup>2</sup> |
| Anidulafungin | 99 (96, 100) | 100<sup>a</sup> |  |  |
| Amphotericin | 100<sup>a</sup> | 96 (92, 100) | 3<sup>b</sup> |  |
| Voriconazole | 90 (83, 97) | 97 (93, 100) | 2<sup>c</sup> |  |
| Itraconazole | 87 (80, 95) | 99 (95, 100) | 1<sup>d</sup> | 1<sup>e</sup> |

<sup>a</sup>Concordance was near perfect and was not possible to calculate confidence interval.
<sup>1</sup>Very Major Discrepancy (VMD): <sup>b</sup>two *A. flavus* isolates and one *A. terreus* isolate were categorised as sensitive to amphotericin by Micronaut whereas CLSI classified them as resistant.
<sup>c</sup>Two different isolates of *A. fumigatus* were categorised as sensitive to voriconazole by MN but CLSI classified them as resistant.
<sup>d</sup>One isolate of *A. fumigatus* was also classified as wild type by MN whilst CLSI classified as non-wild type
<sup>2</sup>Major discrepancy (MD): <sup>e</sup>one isolate of *A. fumigatus* was classified as resistant to Itraconazole by MN but was sensitive in CLSI.

In total nine isolates showed major and very major discrepancies and upon repeating the test, six isolates still gave discrepant results. For the very major discrepancies, two isolates of *A flavus* and one isolate of *A terreus* had MIC of 1mg/L compared to MIC of 2mg/L in CLSI method for isolates. In addition, two isolates of A fumigatus gave voriconazole MICs of 0.5mg/L and 0.25mg/L whereas CLSI MICs were 2mg/L for both isolates. As for major discrepancies only 1 isolate of *A fumigatus* has Itraconazole MICs of 4mg/L compared to CLSI MIC of 0.25mg/L (table 2).

Essential agreement amongst different experiments and observers was very high (≥ 90%); results are shown in table 3. Interexperimental essential agreement was 98%, 99%, 94% and 92% for Amphotericin, Anidulafungin, Itraconazole and Voriconazole respectively. Interobserver agreement was 96%, 95%, 94% and 97% for Amphotericin, Anidulafungin, Itraconazole and Voriconazole respectively.

**Table 3:** Essential reproducibility of Micronaut-AM method as determined by percent inter-experimental and percent inter-observer agreement.

| Drug | % Interexperimental agreement | % Inter-observer agreement |
| --- | --- | --- |
| Amphotericin | 98 | 96 |
| Anidulafungin | 99 | 95 |
| Itraconazole | 94 | 94 |
| Voriconazole | 92 | 97 |

## Discussion

Despite increasing *Aspergillus species* resistance to antifungal and associated treatment failure, few laboratories in the UK have access to susceptibility testing.^16^ In a survey on mycology diagnostic capacity in 2017 only 17% of Laboratories had access to local susceptibility testing.^9^ Availability of a robust commercial *Aspergillus species* susceptibility testing method would help overcome this challenge. We have presented here the evaluation and validation of a colorimetric method, Micronaut-AM, for Aspergillus susceptibility testing.

Micronaut-AM (MN) showed significant concordance (P <0.0001) with the reference CLSI broth microdilution method and generated minimum inhibitory concentrations (MICs) and minimum effective concentrations (MECs) that were very close and comparable to those obtained by reference method. In addition, MN reliably detected azole resistant aspergillus isolates and showed very good interobserver and interexperimental reproducibility. Furthermore, MICs and MECs end point were easily determined as consistently indicated by first blue and first purple colour respectively.

MN showed high concordance with CLSI for isolates and antifungals tested and overall essential agreement was ≥90% for anidulafungin, amphotericin and voriconazole. Itraconazole gave slightly lower essential agreement of 87%. These results are similar to those obtained by other researchers in the evaluation of Sensititre YeastOne colorimetric method.^10,17,18^ Categorical agreement and essential interexperimental and interobserver agreements were equally very high and MN showed percent agreement greater than 90% for all drugs tested with very few major and very major discrepancies.

Of the very major discrepancies, Amphotericin had the highest discrepancy of three, where isolates were classified by MN as wild-type (WT) but as non-WT by CLSI. Voriconazole showed very major discrepancy of two where itraconazole had just one very major discrepancy. Since MN is based on EUCAST these discrepancies may be explained partly by the difference between EUCAST and CLSI methods, which include higher glucose content (2%) RPMI broth and the use of flat-bottom microdilution plate in EUCAST method. Similar discrepancies were reported by Pfaller et al.,^19^ when comparing EUCAST and CLSI and Wang et al.,^17^ when comparing YO with CLSI.

Overall MN reliably identified azole resistance. Almost all isolate that were resistant were identified by MN method and only one major discrepancy was observed, where an isolate classified as resistant by CLSI gave wild type MIC. Interestingly this isolate was also classified as wild type by azole resistance screening kit, VIPcheck™ system.^20^ This may be due to the isolate losing resistance during passaging.

For echinocandins MEC was consistently indicated by the first purple well when plates were read between 18 and 24h incubations; after 48h incubation all wells were pink. This was confirmed by viewing aberrant hyphal growth under inverted microscope both at 24h and 48h incubation (data not shown). In this method, echinocandin susceptibility results can be reliably reported within 24 incubation and we found that 10^4^ CFU was optimal inoculum as reported for YO by Siop et al.^15^ However, our concordance was higher than those reported by these researchers, especially anidulafungin, which is probably due to the optimised almarBlue concentration.

In conclusion, our study has clinical implications to improve local fungal diagnostics where routine microbiology laboratories can provide timely access to *Aspergillus species* susceptibility testing. Micronaut-AM was robust and easy to use and showed good agreement with reference broth micro-dilution method results for all antifungal agents tested. In addition, echinocandin susceptibility results were available within 24 hours incubation and this method reliably detected azole resistance. This colorimetric method is very promising and appears to be a suitable alternative susceptibility testing to labour intensive reference broth microdilution methods for *Aspergillus species*. Therefore, further multicentre evaluation and extension to other moulds should be performed to confirm these results.

### Limitations of this study

Three limitations of this study are that the two methods, MN and CLSI were not run concomitantly; resistance organisms were not confirmed genotypically and did not include echinocandin resistance.

## Acknowledgement

Authors would like to thank Winston BANYA who assisted with statistical analysis, Dr Richard BARTON who reviewed the manuscript and Bioconnections for providing the kit. We also appreciate Dr Alireza ABDOLRASOULI for his advice on methodology.

## References

1. Kosmidis, C. & Denning, D. W. The clinical spectrum of pulmonary aspergillosis. Thorax 70, 270–277 (2015).

2. Gregg, KS and Kauffman, C. Invasive Aspergillosis: Epidemiology, Clinical Aspects, and Treatment. Semin. Respir. Crit. Care Med. 36, 662–672 (2015).

3. Taccone, F. S. et al. Epidemiology of invasive aspergillosis in critically ill patients: clinical presentation, underlying conditions, and outcomes. Crit. Care 19, 7 (2015).

4. Schelenz, S. et al. British Society for Medical Mycology best practice recommendations for the diagnosis of serious fungal diseases. Lancet Infect. Dis. 15, 461–474 (2015).

5. Lestrade, P. P. A., Meis, J. F., Melchers, W. J. G. & Verweij, P. E. Triazole resistance in Aspergillus fumigatus: recent insights and challenges for patient management. Clin. Microbiol. Infect. 25, 799–806 (2019).

6. Beer, K. D. et al. Multidrug-Resistant Aspergillus fumigatus Carrying Mutations Linked to Environmental Fungicide Exposure — Three States, 2010–2017. MMWR. Morb. Mortal. Wkly. Rep. 67, 1064–1067 (2018).

7. CLSI. (2017). Available at: https://clsi.org/search/?q=mould. (Accessed: 24th August 2019)

8. EUCAST. (2017). Available at: http://www.eucast.org/astoffungi/methodsinantifungalsusceptibilitytesting/susceptibility_testing_of_moulds/. (Accessed: 23rd August 2019)

9. Schelenz, S. et al. National mycology laboratory diagnostic capacity for invasive fungal diseases in 2017: Evidence of sub-optimal practice. J. Infect. 79, 167–1731. Schelenz S, Owens K, Guy R, et al. Natio (2019).

10. Mello, E. et al. Susceptibility Testing of Common and Uncommon Aspergillus Species against Posaconazole and Other Mold-Active Antifungal Azoles Using the Sensititre Method. Antimicrob. Agents Chemother. 61, (2017).

11. Castro, C., Mc, S., Flores, B., Espinel-Ingroff, A. & Martín-Mazuelos, E. Comparison of the Sensititre YeastOne colorimetric antifungal panel with a modified NCCLS M38-A method to determine the activity of voriconazole against clinical isolates of Aspergillus spp. J. Clin. Microbiol. 42, 4358–4360 (2004).

12. García-Martos, P. et al. [In vitro activity of amphotericin B, itraconazole and voriconazole against 20 species of Aspergillus using the Sensititre microdilution method]. Enferm. Infecc. Microbiol. Clin. 23, 15–18 (2005).

13. Meletiadis, J., Mouton, J. W., Meis, J. F. G. M., Bouman, B. A. & Verweij, P. E. Comparison of the Etest and the sensititre colorimetric methods with the NCCLS proposed standard for antifungal susceptibility testing of Aspergillus species. J. Clin. Microbiol. 40, 2876–2885 (2002).

14. Dunne K, Renwick J, McElvaney NG, Chotirmall S, Meis JF, Klaassen CHW, Murphy P, R. T. Aspergillus fumigatus in an Irish Cystic Fibrosis Patient Cohort◻; Epidemiology and Antifungal SuscetibilityÂ | Aspergillus & Aspergillosis Website. Available at: https://www.aspergillus.org.uk/content/aspergillus-fumigatus-irish-cystic-fibrosis-patient-cohort-epidemiology-and-antifungal. (Accessed: 26th August 2019)

15. Siopi, M., Pournaras, S. & Meletiadis, J. Comparative Evaluation of Sensititre YeastOne and CLSI M38-A2 Reference Method for Antifungal Susceptibility Testing of Aspergillus spp. against Echinocandins. J. Clin. Microbiol. 55, 1714–1719 (2017).

16. Bueid, A. et al. Azole antifungal resistance in Aspergillus fumigatus: 2008 and 2009. J. Antimicrob. Chemother. 65, 2116–2118 (2010).

17. Wang, H. C., Hsieh, M. I., Choi, P. C. & Wu, C. J. Comparison of the sensititre YeastOne and CLSI M38-A2 microdilution methods in determining the activity of amphotericin B, itraconazole, voriconazole, and posaconazole against aspergillus species. J. Clin. Microbiol. 56, (2018).

18. Linares, M. J. et al. Susceptibility of filamentous fungi to voriconazole tested by two microdilution methods. J. Clin. Microbiol. 43, 250–253 (2005).

19. Pfaller, M. et al. Comparison of the broth microdilution methods of the European Committee on Antimicrobial Susceptibility Testing and the Clinical and Laboratory Standards Institute for testing itraconazole, posaconazole, and voriconazole against Aspergillus isolates. J. Clin. Microbiol. 49, 1110–1112 (2011).

20. VIPcheckTM system, Netherlands. (2019). Available at: https://www.vipcheck.nl. (Accessed: 25th August 2019)

